# A Multidimensional Approach to Understanding the Emergence of Sex Differences in Internalizing Symptoms in Adolescence

**DOI:** 10.1101/2022.05.05.490817

**Authors:** Bianca Serio, Robert Kohler, Fengdan Ye, Sarah D Lichenstein, Sarah W Yip

## Abstract

Women are more vulnerable to internalizing disorders (e.g., depression and anxiety). This study took an integrative, developmental approach to investigate multidimensional factors associated with the emergence of sex differences in internalizing symptoms, using data from the Adolescent Brain Cognitive Development (ABCD) study. Indices of sex hormone levels (dehydroepiandrosterone, testosterone, and estradiol), physical pubertal development, task-based functional brain activity, family conflict, and internalizing symptoms were drawn from the ABCD study’s baseline sample (9-to 10-year-old; *N* = 11,844). Principal component analysis served as a data-driven dimensionality reduction technique on the internalizing subscales to yield a single robust measure of internalizing symptoms. Moderated mediation analyses assessed whether associations between known risk factors and internalizing symptoms vary by sex. Results revealed direct and indirect effects of physical pubertal development on internalizing symptoms through family conflict across sexes. No effects were found of sex hormone levels or amygdala response to fearful faces on internalizing symptoms. Females did not report overall greater internalizing symptoms relative to males, suggesting that internalizing symptoms have not yet begun to increase in females at this age. Findings provide an essential baseline for future longitudinal research on the endocrine, neurocognitive, and psychosocial factors associated with sex differences in internalizing symptoms.

## 1. Introduction

Internalizing disorders—e.g., depression and anxiety—are a significant form of psychopathology that manifests during adolescence [1, 2] and disproportionally affects women relative to men [3] [4]. Such sex differences in internalizing symptoms have been recorded as early as age 9 [5], although this estimate significantly varies between studies [e.g., 6, 7-9]. Studying the emergence of sex differences in internalizing symptoms from a developmental perspective and prior to peak prevalence may help to identify when sex differences in internalizing symptomatology emerge, as well as neurodevelopmental mechanisms placing females at higher risk. Given that adolescence is marked by drastic changes across multiple domains, we take a multidimensional approach to test a novel theoretical model of biological and psychosocial factors contributing to the emergence of internalizing disorders in youth, including sex hormone levels, brain function, pubertal development, and family conflict.

The emergence of differences in internalizing symptoms appears tightly linked with the onset of puberty [10]. Among females, more advanced pubertal development is associated with greater depression and anxiety even when controlling for age [11, 12]. Puberty is driven by a steep increase in circulating sex steroid hormones, namely dehydroepiandrosterone (DHEA), testosterone, and estradiol [13]. Although there is currently no consensus on the direction of relationships between internalizing symptoms and sex hormone levels, rises in DHEA [14, 15], testosterone [16, 17], and estradiol [16, 18] have all been associated with increased internalizing symptoms during puberty, particularly among females.

Rising sex hormone levels during adolescence also drive brain maturation, particularly the development of affective neural circuits associated with internalizing symptoms [19]. Animal research suggests that estrogen receptors in the amygdala mediate the effects of estradiol on anxiety-like behaviors [20]. This preclinical work is supported by a growing body of human neuroimaging research linking the effects of DHEA [21], testosterone [22], and estradiol [23] to the amygdala’s response to emotional stimuli.

Relative to adults, adolescents show increased amygdala reactivity to socio-emotional cues such as emotional faces [24]. The adolescent brain is also particularly sensitive to cues from the family environment [25]. Evidence suggests that the effects of negative maternal parenting and aggression on adolescents’ depressive symptoms are mediated by amygdala resting-state functional connectivity to frontal [26] and temporal [27] regions respectively. Patterns of amygdala activity and connectivity may thus be a pathway through which adolescents with negative family environments develop internalizing disorders.

Pubertal development also interacts with psychosocial factors that may influence the development of internalizing symptoms. Much of the extant literature focuses on pubertal timing and suggests that adolescents with early pubertal timing are more likely to develop internalizing symptoms throughout adolescence [28] and adulthood [29], particularly females [30]. Furthermore, pubertal timing is associated with internalizing symptoms—both cross-sectionally and prospectively—when pubertal development is measured by physical assessments, but not by sex hormone levels [31]. Although this lack of significant findings for sex hormones could be partly explained by their high interpersonal variability [32], associations between pubertal development and internalizing symptoms may be more meaningfully explained by psychosocial mechanisms rather than biological mechanisms. For instance, the family environment strongly influences adolescent wellbeing, as underpinned by self-esteem, in different ways [33]. Family cohesion may buffer the association between internalizing behaviors and peer victimization [34], a factor affecting self-esteem in females during puberty [35], and parental support may protect against maladjustment to peer-victimization, especially in females [36]. Conversely, harsh parenting may exacerbate the association between pubertal timing and internalizing symptoms in adolescent females [37]. Therefore, family dynamics may interact with psychosocial stress during pubertal development, influencing internalizing symptoms and these relationships may vary between male and female adolescents.

In sum, endocrine, neurocognitive, and psychosocial factors associated with sex differences in internalizing symptoms tightly interact with one another and cannot be fully understood in isolation, particularly during adolescence, when these factors are dynamically changing. Despite observed associations between internalizing symptoms and rising sex hormone levels, affective neurodevelopment, and changing family dynamics, research is needed assessing all these risk factors concurrently in the context of emerging sex differences [38]. Furthermore, these relationships have mostly been observed in small and homogeneous samples. This study thus uses a large-scale heterogeneous dataset to investigate the putative interacting contributions of endocrine, neurocognitive, and psychosocial factors on emerging sex differences in internalizing symptoms in adolescence, as proposed in our theoretical model (Figure 1).

**Figure 1.**
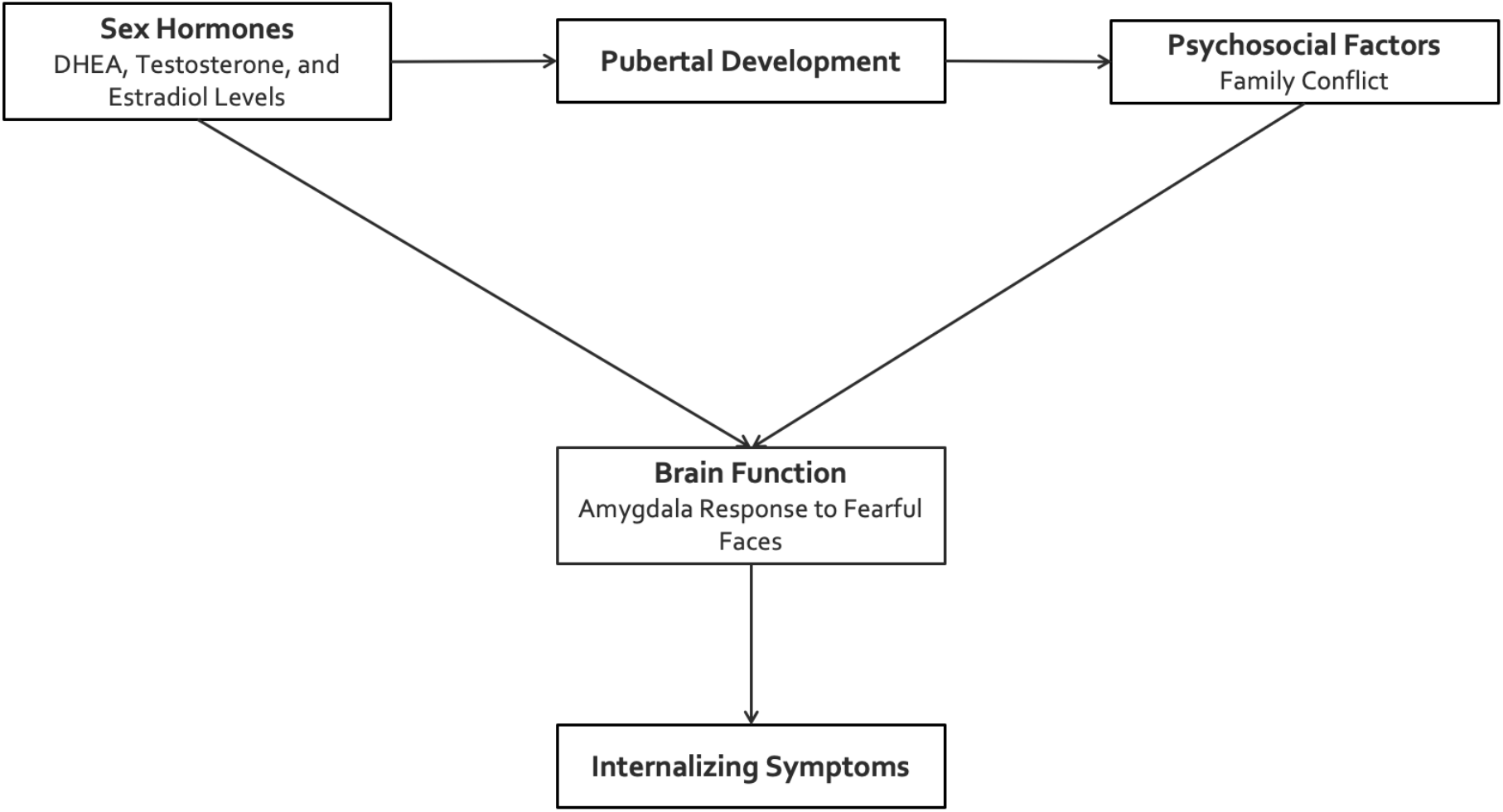
Proposed theoretical model. *Note*. DHEA, dehydroepiandrosterone. This model suggests that sex hormones (i.e., DHEA, testosterone, and estradiol) drive pubertal development; pubertal development influences psychosocial factors (i.e., family conflict); sex hormones and psychosocial factors lead to internalizing symptoms through brain function (i.e., amygdala response to fearful faces), the putative neural substrate of internalizing symptoms.

This study tests our novel theoretical model in a large and diverse sample of 9- to 10- year-olds. We used heterogeneous, large-scale data from the Adolescent Brain Cognitive Development (ABCD) study, including indices of sex hormone levels, physical pubertal development, task-based functional brain activity, and family conflict. Studying 9- to 10-year-olds uniquely captures the transition from childhood to adolescence and thus the emergence of sex differences in internalizing symptoms, prior to peak onset. It therefore also enables the establishment of an empirical baseline from which relationships between endocrine, neurocognitive, and psychosocial factors may be assessed relative to internalizing symptoms in later data releases of the ABCD study. To test our proposed theoretical model, we hypothesized direct and indirect effects of (1) sex hormone levels (i.e., DHEA, testosterone, and estradiol) on internalizing symptoms through amygdala response to fearful faces; (2) family conflict on internalizing symptoms through amygdala response to fearful faces; and (3) physical pubertal development on internalizing symptoms through family conflict. These effects were hypothesized across sexes and expected to be stronger in females relative to males.

## 2. Methods

### 2.1. Participants

The ABCD study is the largest longitudinal consortium study of brain development in the United States [39], comprehensively measuring wide-ranging factors affecting adolescent neurodevelopment and health [40]. Supported by the National Institute of Mental Health (NIMH), this study covers the transition from childhood to adulthood by prospectively following a baseline cohort of 9-to 10-year-old youth for ten years [41]. In this study, we used the ABCD sample at baseline (Release 3.0), including 11,844 9-to 10-year-olds (M_age_ = 118.97 ± 7.50 months) – see Table 1 for demographics. A detailed account of recruitment considerations and procedures is summarized in [42, 43]. Written parent consent and verbal child assent was required prior to participation [44].

**Table 1.** Demographic characteristics of the sample

| Variable | *n* | % |
| --- | --- | --- |
| **Sex** |  |  |
| Male | 6,182 | 52.20 |
| Female | 5,662 | 47.80 |
| **Ethnicity** |  |  |
| Hispanic/Latinx | 2,405 | 20.31 |
| Non-Hispanic/Latinx | 9,286 | 78.40 |
| Unknown | 153 | 1.29 |
| **Race** |  |  |
| White | 7,507 | 63.38 |
| Black | 1,860 | 15.70 |
| Asian | 274 | 2.31 |
| Mixed | 2,041 | 17.23 |
| Unknown | 162 | 1.37 |
| **Total combined household income (past 12 months)** |  |  |
| < \$5,000 | 416 | 3.51 |
| \$5,000 – \$11,999 | 419 | 3.54 |
| \$12,000 – \$15,999 | 274 | 2.31 |
| \$16,000 – \$24,999 | 522 | 4.41 |
| \$25,000 – \$34,999 | 653 | 5.51 |
| \$35,000 – \$49,999 | 932 | 7.87 |
| \$50,000 – \$74,999 | 1,498 | 12.65 |
| \$75,000 – \$99,999 | 1,568 | 13.24 |
| \$100,000 – \$199,999 | 3,308 | 27.93 |
| > \$200,000 | 1,238 | 10.45 |
| Unknown | 1,016 | 8.58 |

*Note.* *n*, number of subjects. Given that sample sizes for our different analyses varied according to data availability, the demographic characteristics reported here relate to all subjects included at least once in our analyses.

### 2.2. Measures

#### 2.2.1. Internalizing Symptoms

Internalizing symptoms were measured with the parent/caregiver-report version of the Child Behavior Checklist (CBCL) for 6- to 18-year-old youth, a standardized questionnaire broadly assessing youth psychopathology on dimensional scales [45]. The CBCL includes 134 items contributing to five empirically-derived dimensions (i.e., internalizing, externalizing, thought, attention, and social symptoms), each item describing emotional and behavioral problems on 3-point Likert scales. Seven CBCL internalizing symptom subscales were selected for the current analyses, i.e., Anxious/Depressed, Withdrawn/Depressed, Somatic, DSM Depressed, DSM Anxiety, DSM Somatic, and Stress subscales. Analyses were conducted on norm-referenced t-scores (M = 50, SD = 10).

#### 2.2.2. Sex Hormone Levels

The ABCD study sampled DHEA and testosterone in both males and females, and 17-b estradiol in females only. Research assistants collected salivary samples via passive drool [46]. The ABCD study saliva collection protocol and sex hormone assay specifications are summarized by Herting, Uban [47]. Inclusion criteria for the hormonal data were based on a decision tree adapted from Herting, Uban [47] (Figure 2), summarizing the steps taken to conduct a quality control assessment of each replicate in order to establish a single final estimate for DHEA (*n* = 10,932) and testosterone (*n* = 10,978) levels in both sexes, and estradiol (*n* = 5,119) levels in females only.

**Figure 2.**
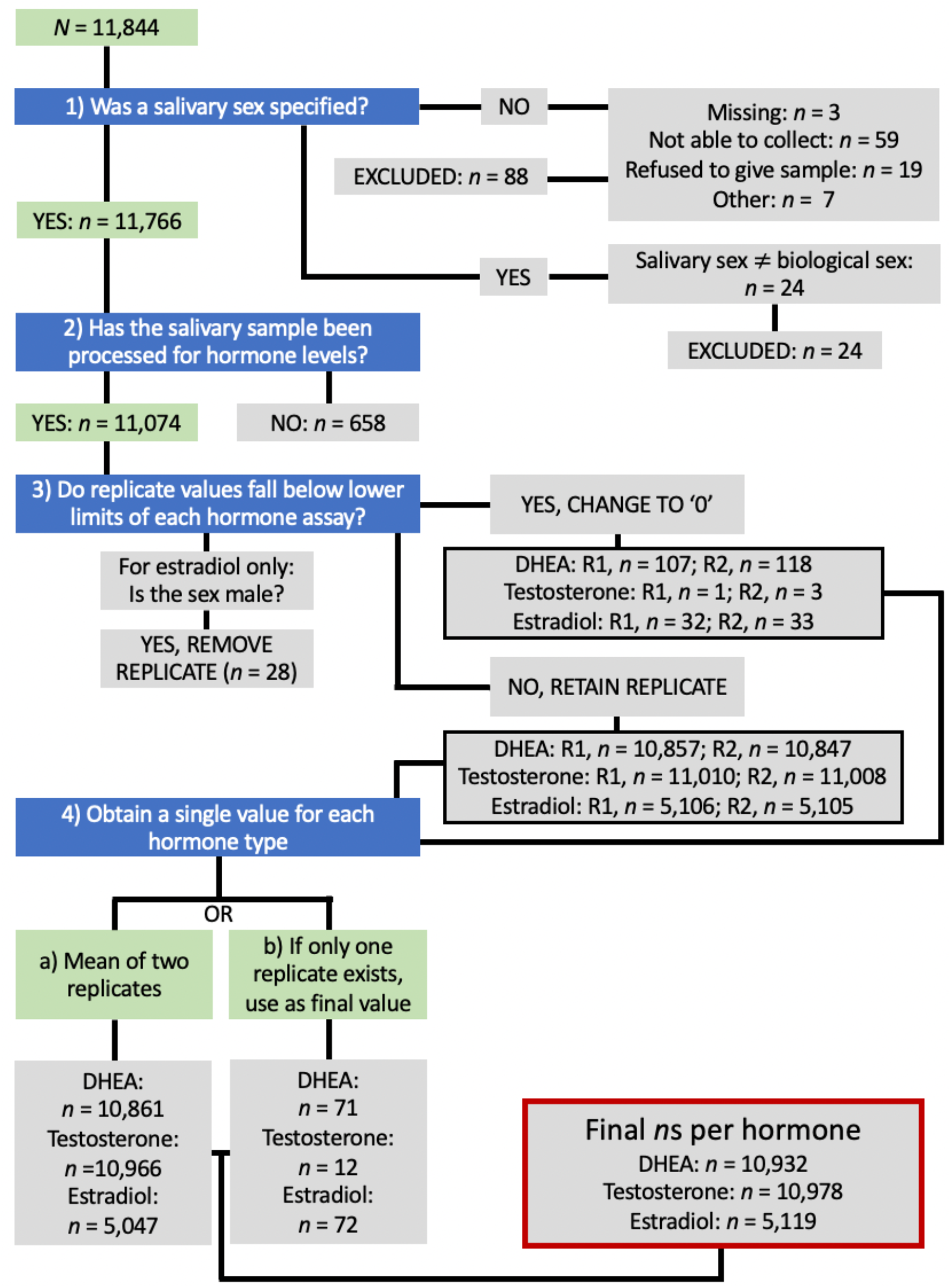
Decision tree for the quality control inclusion criteria of sex hormone levels. *Note. n*, number of subjects; R1, replicate 1; R2, replicate 2. Decision tree adapted from Herting, Uban [47]. Data was retained if (1) salivary sex was specified and matched the biological sex; (2) the salivary sample was processed for hormone levels, i.e., at least one replicate for at least one of the three hormones was processed; (3) replicates falling below lower detection limits were valued as 0 pg/ml, and estradiol replicates measured in males were removed; (4) a final value for each hormone type was obtained by: a) calculating the mean of the two replicates; or b) if only one replicate was available, using it as the final value.

#### 2.2.3. Physical Pubertal Development

Physical pubertal development was measured with the Pubertal Development Scale (PDS), a widely used and standardized questionnaire assessing physical pubertal development [48]. It includes five items in total, assessing perceived changes in: body hair, skin, and height (both sexes); voice deepening and facial hair (males only); and breast development and menarche (females only). All items are scored on 4-point Likert scales (with higher scores indicating more advanced pubertal development), except menarche (for which 1 indicates “menstruation has not yet begun” and 4 indicates “menstruation has begun”). PDS scores were computed by averaging item scores, yielding a final score out of four. The PDS was administered to both youth and parents, but we only considered parental reports given their greater reliability and internal consistency on the PDS compared to adolescent self-reports [49].

#### 2.2.4. Family Conflict

Family conflict was measured using the Family Environment Scale (FES), a self-report questionnaire assessing the general social climate of families [50]. The questionnaire consists of nine true or false statements scored with 0 or 1. These scores were summed to yield a final score out of nine, with higher scores indicating greater family conflict.

#### 2.2.5. fMRI Emotional Face N-Back Task

Emotional reactivity was measured with the fMRI emotional face n-back task, a validated experimental paradigm assessing emotional reactivity and working memory [51, 52], previously associated with the engagement of the fronto-amygdala circuitry [53]. It includes both low and high memory load conditions (0-back and 2-back conditions respectively), for which subjects are required to indicate whether a given stimulus is a “match” or a “no match” to the stimulus presented at the beginning of the block (0-back condition) or two trials back (2-back condition). The task comprises two runs of eight blocks, each run consisting of four 0-back and four 2-back condition blocks. The stimuli included pictures of happy, fearful, and neutral faces, as well as places [54, 55]. Each block, for both 0-back and 2-back conditions, included 10 trials. In order to assess emotional reactivity to fearful faces, the amygdala was selected as the region of interest, as measured by mean beta weights for the fearful versus neutral faces contrast. Data from the fMRI emotional face n-back task were included (*n* = 7,712) based on the recommended quality control inclusion criteria provided in ABCD Release 3.0. These criteria and the imaging procedure are outlined in the Supplementary Material.

### 2.3. Analytic Plan

The following analyses were conducted on the ABCD study’s Curated Annual Release 3.0, made publicly available on November 11th, 2020, and downloaded on February 18th, 2021, from the NIMH Data Archive (https://nda.nih.gov/abcd/query/abcd-annual-releases.html). Subjects with missing values, or whose data did not meet the quality control inclusion criteria for sex hormone levels and the fMRI Emotional Face N-Back Task (see sections 2.2.2. and 2.2.5.) were excluded within the variables’ respective data files. All statistical analyses were conducted in R (version 4.0.3).

#### 2.3.1 Descriptive Statistics and Sex Differences

Descriptive statistics were computed for all variables of interest (i.e., CBCL scores on the internalizing symptom subscales; sex hormone levels; physical pubertal development; family conflict; and amygdala response to fearful faces) and, given that all data violated the normality assumption and were mostly measured on ordinal scales, sex differences were assessed with the non-parametric Mann-Whitney U test and effect sizes were measured with rank-biserial correlation (*r*_rb_) [56]. Bonferroni correction was used to reduce the risk of type I errors by accounting for multiple comparisons across all analyses, setting the significance level at .0042 (.05/12).

#### 2.3.2. Principal Component Analysis (PCA)

PCA was performed on the seven CBCL internalizing symptom subscales as a data-driven dimensionality reduction technique which parses variance by yielding summary indices, named principal components, that are orthogonal to one another and in decreasing order of variance explained. It was decided a priori that only the first principal component (PC1) would be retained as a single composite measure of internalizing symptoms to test our hypotheses. Prior to conducting PCA, we assessed sampling adequacy of the data with the Kaiser-Meyer-Olkin (KMO) test (KMO = .76), which measures the proportion of variance among factors that may be common variance and may thus represent latent factors. Suitability for data reduction was also measured with Bartlett’s test of sphericity (χ^2^(21) = 82,637.59, *p* < .001) by testing the null hypothesis that a correlation matrix is an identity matrix, which would imply that variables are unrelated and thus unsuitable for pattern detection. The function prcomp (stats package, R) was used to conduct PCA on the CBCL internalizing symptom subscales. All data were mean-centered and scaled.

#### 2.3.3. Mediation and Moderated Mediation Analyses

Moderated mediation analyses (mediation package, R) were used to test the hypothesized direct and indirect effects of: (1) sex hormone levels (i.e., DHEA and testosterone) on internalizing symptoms through amygdala response to fearful faces; (2) family conflict on internalizing symptoms through amygdala response to fearful faces; and (3) physical pubertal development on internalizing symptoms through family conflict. The direct and indirect effects of estradiol on internalizing symptoms through amygdala response to fearful faces were tested with a simple mediation analysis given that estradiol levels were only assayed in females and thus no moderation by sex could be assessed. Moderated mediation models were evaluated by first testing for an indirect effect through mediation analysis (i.e., effect of the independent variable on the dependent variable through the mediator variable), followed by testing whether sex moderates this effect (see Table 2 for a specification of the variables used to test each hypothesized model).

**Table 2.** Specification of the variables used to test each hypothesized model

| IV | DV | MedV | ModV |
| --- | --- | --- | --- |
| DHEA Levels | Internalizing Symptoms | Amygdala Response to Fearful Faces | Sex |
| Testosterone Levels | Internalizing Symptoms | Amygdala Response to Fearful Faces | Sex |
| Estradiol Levels | Internalizing Symptoms | Amygdala Response to Fearful Faces | - |
| Family Conflict | Internalizing Symptoms | Amygdala Response to Fearful Faces | Sex |
| Pubertal Development | Internalizing Symptoms | Family Conflict | Sex |

*Note.* IV, independent variable; DV, dependent variable; MedV, mediator variable; ModV, moderator variable; DHEA, dehydroepiandrosterone.

Conducting mediation analysis with the mediation R package requires fitting two linear regression models [57]. Model 1 specifies the effects of the independent (and moderator) variables on the mediator variable. Model 2 specifies the effects of the independent, mediator (and moderator) variables on the dependent variable. To proceed with testing for an indirect effect through mediation analysis, a statistically significant effect of the independent variable on the mediating variable (Model 1) must be observed, as well as a statistically significant effect of the mediating variable on the dependent variable when the independent and mediating variables are also included in the model as predictors (Model 2) [58]. Models 1 and 2 were tested for all hypotheses, including sex as the moderator variable (except when testing estradiol effects). All variables were mean-centered and scaled. Given the ABCD data’s nested structure [59], we used the R function lmer (lmerTest package, R) to run generalized linear mixed-effect (GLME) models, which included random nested effects for site and family. To control for potentially confounding effects in models including hormone data, additional covariates – which have been found to influence sex hormone levels in the ABCD study– were included (i.e., collection duration (in minutes) for DHEA, testosterone, and estradiol; time of the day at sample collection (in minutes) for testosterone and estradiol; and caffeine in the past 12h (yes/no) for estradiol) [47].

For hypotheses demonstrating statistically significant effects in Models 1 and 2, moderated mediation was tested in two steps. First, mediation was tested with the function mediate (mediation package, R) separately in males and females. Mediation was conducted with 2,000 iterations of bias-corrected and accelerated (BCa) bootstrap confidence intervals [60] to provide more accurate estimates and correct for deviations from normality [61]. The effect size (*R*^2^) for the mediator on the dependent variable was obtained with the function mediation (MBESS package, R). Next, moderation was tested with the function test.modmed (mediation package, R) including sex as the moderator, also with 2,000 iterations of BCa bootstrap confidence intervals.

## 3. Results

### 3.1. Descriptive Statistics and Sex Differences

Statistical results for all between-sex comparisons are shown in Table 3. Females did not show higher overall internalizing symptoms relative to males. Males displayed higher mean scores on the Withdrawn/Depressed (|*μ*_Difference_| = 1.54; *U* = 15,551,276; *p* < .001), DSM Depression (|*μ*_Difference_| = 0.67; *U* = 16,371,052, *p* < .001), DSM Anxiety (|*μ*_Difference_| = 0.86; *U* = 14,094,259, *p* < .001), and Stress (|*μ*_Difference_| = ; 1.10; *U* = 15,878,058, *p* < .001) subscales, whereas females only showed higher mean scores on the Somatic subscale (|*μ*_Difference_| = 0.43, *U* = 18,192,098, *p* < .001). For all other variables of interest, except amygdala response to fearful faces, statistically significant sex differences were found. Specifically, females showed higher mean DHEA levels (|*μ*_Difference_| = 18.13 pg/ml; *U* = 18,153,248, *p* < .001), mean testosterone levels (|*μ*_Difference_| = 3.95 pg/ml, *U* = 17,222,647, *p* < .001), pubertal development (|*μ*_Difference_| = 0.39, *U* = 23,328,564, *p* < .001), and lower family conflict (|*μ*_Difference_| = 0.23, *U* = 16,125,347, *p* < .001), relative to males.

**Table 3.** Mann-Whitney U test statistics for sex differences in scores on all variables of interest

| Variable | Sex |  | U | r<sub>rb</sub> |
| --- | --- | --- | --- | --- |
|  | Males (Mean ± SD) | Females (Mean ± SD) |  |  |
| CBCL Subscales (n = 11,836) |  |  |  |  |
| Anxious/Depressed | 53.81 ± 6.16 | 53.12 ± 5.71 | 17,005,644 | .03 |
| Withdrawn/Depressed | 54.25 ± 6.33 | 52.71 ± 5.03 | 15,551,276* | .11 |
| Somatic | 54.67 ± 5.99 | 55.10 ± 6.09 | 18,192,098* | .04 |
| DSM Depression | 53.92 ± 5.97 | 53.25 ± 5.43 | 16,371,052* | .06 |
| DSM Anxiety | 53.91 ± 6.23 | 53.05 ± 5.98 | 14,094,259* | .19 |
| DSM Somatic | 55.32 ± 6.61 | 55.61 ± 6.62 | 17,501,550 | .00 |
| Stress | 53.85 ± 6.48 | 52.75 ± 5.38 | 15,878,058* | .09 |
| Sex Hormone Levels (pg/ml) |  |  |  |  |
| DHEA (n = 10,932) | 54.92 ± 41.91 | 73.05 ± 54.96 | 18,153,248* | .22 |
| Testosterone (n = 10,978) | 32.09 ± 19.93 | 36.04 ± 18.10 | 17,222,647* | .15 |
| Estradiol (n = 5,119) | - | 1.04 ± 0.50 | - | - |
| PDS score (n = 11,372) | 1.18 ± 0.35 | 1.57 ± 0.56 | 23,328,564* | .45 |
| FES score (n = 11,817) | 2.16 ± 1.97 | 1.93 ± 1.93 | 16,125,347* | .07 |
| Amygdala response to fearful faces<br>(n = 7,712) | 0.05 ± 0.49 | 0.06 ± 0.44 | 7,407,891 | .00 |

Note. SD, standard deviation; $df$ , degrees of freedom; U, Mann-Whitney U test statistic; $r_{rb}$ , rank-biserial correlation; $n$ , number of subjects; CBCL, Child Behavior Checklist; DSM, Diagnostic and Statistical Manual of Mental Disorders; DHEA, dehydroepiandrosterone; PDS, Pubertal Development Scale; FES, Family Environment Scale. Bold denotes statistically significant sex differences at $p < .01$ ; \* denotes statistically significant sex differences at the Bonferroni corrected significance level ( $p < .0042$ ).

The emotional face n-back task’s validity in adequately engaging the amygdala was confirmed with a one-sample t-test, showing that the beta-weight for the contrast between fearful and neutral faces in the amygdala was significantly different from 0, *t*(7,711) = 9.58, *p* < .001.

### 3.2. Principal Component Analysis (PCA)

PCA yielded a first principal component (PC1) that accounted for 64.09% of the variance in all the CBCL internalizing symptom subscales. The Stress subscale contributed the most to PC1 (17.72%), followed by the Anxious/Depressed (17.24%), DSM Anxiety (16.90%), DSM Depression (15.82%), Withdrawn/Depressed (12.45%), Somatic (11.39%), and DSM Somatic (8.46%) subscales. The variance explained by each principal component as well as the factor loadings, showing the contributions of each factor to each principal component, are summarized in the Supplementary Tables S1 and S2.

### 3.3. Mediation and Moderated Mediation Analyses

GLME models did not indicate significant direct or indirect effects of sex hormone levels (i.e., DHEA (*n* = 7,138), testosterone (*n* = 7,157), and estradiol (*n* = 3,420)) on internalizing symptoms through amygdala response to fearful faces. No significant direct and indirect effects of family conflict on internalizing symptoms through amygdala response to fearful faces (*n* = 7,698) were found either. A statistically significant direct effect of family conflict on internalizing symptoms was observed, *t*(6,758) = 5.62, β = 0.089, *p* < .001, but this effect was not moderated by sex (as tested with the family conflict by sex interaction). Given that GLME models testing the effects of sex hormone level and family conflict on internalizing symptoms through amygdala response to fearful faces did not yield the required significance to test for a mediation effect, no additional analyses related to these hypotheses were conducted (full statistical results are available in in Supplementary Table S3).

The GLME model testing direct and indirect effects of physical pubertal development on internalizing symptoms through family conflict (*n* = 11,346) yielded the required significant effects to test for a mediation effect, i.e., effects of pubertal development on family conflict, *t*(1,125) = 2.12, β = 0.026, *p* = .034, and of family conflict on internalizing symptoms, *t*(1,022) = 6.59, β = 0.086, *p* < .001, as well as a statistically significant direct effect of physical pubertal development on internalizing symptoms, *t*(1,122) = 3.75, β = 0.049, *p* < .001, although this effect was not moderated by sex (as tested with the pubertal development by sex interaction; Table 4). Moderated mediation analysis revealed statistically significant indirect effects in both males and females, with an effect size for family conflict on pubertal development of *R*^2^ = .03% (Figure 3; Table 5). However, this indirect effect was not moderated by sex.

**Table 4.** Test statistics of the GLME models required to conduct mediation analysis for Hypothesis 3

| Model | Predictor | b | df | t | p |
| --- | --- | --- | --- | --- | --- |
| 1 | PDS | 0.026 | 1,125 | 2.12 | **.034** |
|  | Sex (M) | 0.133 | 1,112 | 6.46 | **.000\*** |
|  | PDS*Sex (M) | 0.004 | 1,128 | 0.20 | .841 |
| 2 | PDS | 0.049 | 1,122 | 3.75 | **.000\*** |
|  | Sex (M) | 0.162 | 1,132 | 7.92 | **.000\*** |
|  | FES | 0.086 | 1,022 | 6.59 | **.000\*** |
|  | PDS*Sex (M) | 0.025 | 1,121 | 1.15 | .252 |
|  | FES*Sex (M) | -0.005 | 1,013 | -0.30 | .765 |

*Note.* GLME, generalized linear mixed-effect; b, standardized beta coefficient; *df*, degrees of freedom; PDS, Pubertal Development Scale (score); FES, Family Environment Scale (score); M, male. Bold denotes statistical significance at $p < .05$ ; \* denotes statistical significance at $p < .001$ . Model 1 predicts family conflict (FES), and Model 2 predicts internalizing symptoms (PC1 scores).

**Table 5.**
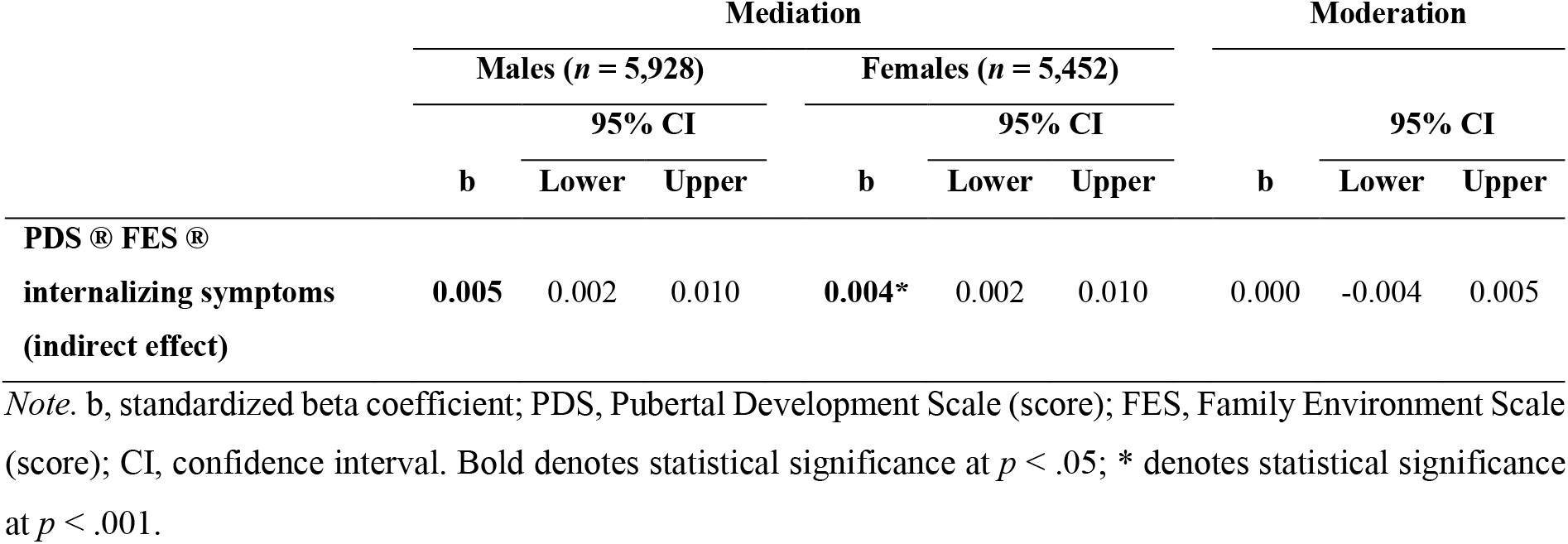
Test statistics of the moderated mediation analysis for Hypothesis 3

**Figure 3.**
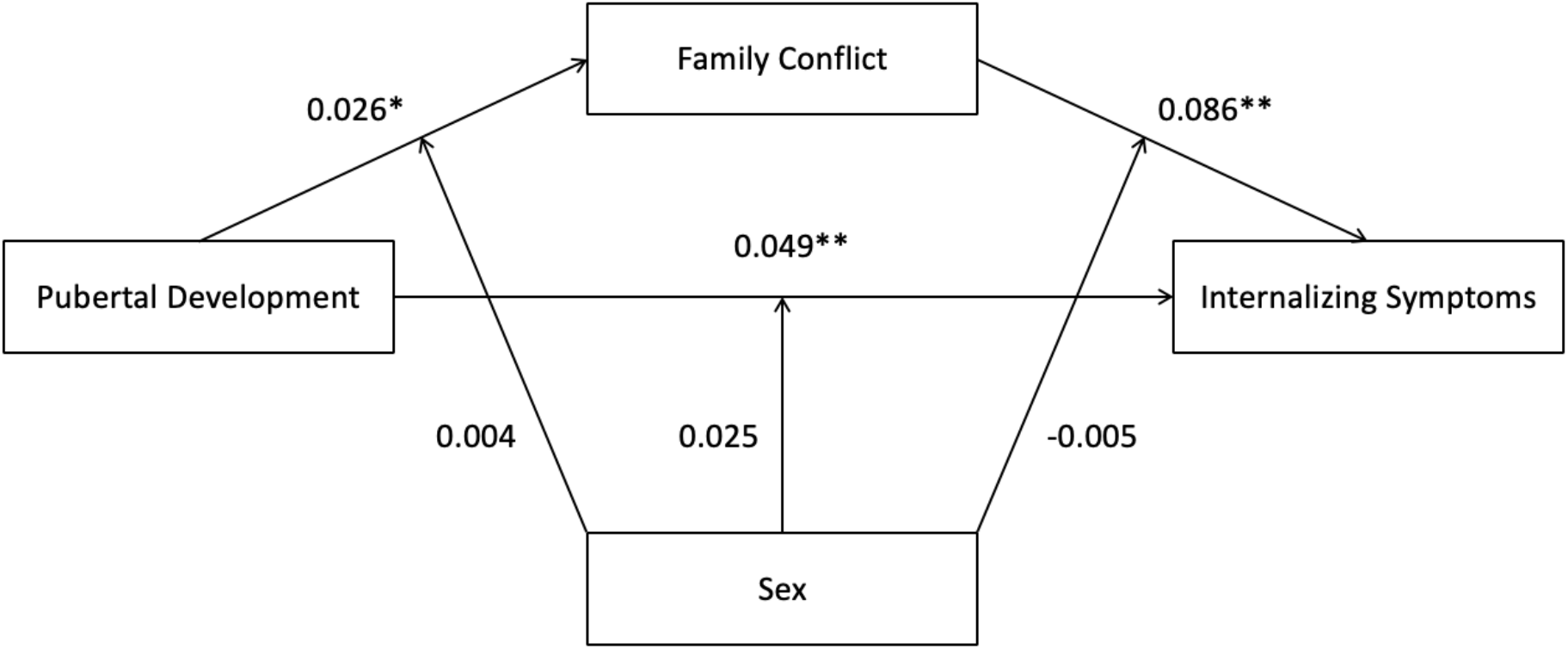
Direct and indirect effects of physical pubertal development on internalizing symptoms through family conflict. *Note*. Reported numbers represent standardized beta coefficients. * denotes statistically significant effects at *p* < .05, ** denotes statistically significant effects at *p* < .001.

## 4. Discussion

This is the largest study to date taking a developmental approach to investigate the mechanisms placing females at greater risk of internalizing psychopathology, including endocrine, neurocognitive, and psychosocial factors, by attempting to capture emerging sex differences in internalizing symptoms. We used heterogeneous data including indices of sex hormone levels, physical pubertal development, task-based functional brain activity, and family conflict to test a novel theoretical model (Figure 1) in the ABCD study’s large and diverse 9-to 10-year-old sample. Mediation analyses indicated significant direct and indirect effects of physical pubertal development on internalizing symptoms through family conflict, although these effects were not moderated by sex. In fact, females did not report overall higher levels of internalizing symptoms. Furthermore, no endocrine or neurocognitive effects were observed. Given that adolescence is a dynamic developmental period, our findings broadly encourage testing our model longitudinally, specifically at later stages of puberty. This first analysis provides an essential empirical baseline from which relationships between endocrine, neurocognitive, and psychosocial factors may be assessed relative to internalizing symptoms in later data releases of the ABCD study.

### 4.1. Greater Prevalence of Internalizing Symptoms in Females is not yet Observable at Ages 9-10

Females in the ABCD sample did not report overall greater internalizing symptoms, scoring higher than males on only one out of seven CBCL subscales (i.e., somatic symptoms – consistent with previous findings [62]). While sex differences in CBCL subscales were modest at this early timepoint, these results nonetheless suggest that future research examining internalizing symptom subtypes may help to explain inconsistencies in previous literature on the presence and directionality of sex differences in preadolescents [5-9]. Prior studies have typically relied on overall scores for internalizing symptoms rather than systematically examining symptom subtypes, which may be a potential reason for mixed results. Therefore, in addition to suggesting that greater prevalence of internalizing symptoms in females may not yet be detectable at ages 9-10, our findings also suggest the importance of considering patterns in symptomatology when studying sex differences in internalizing psychopathology.

### 4.2. Pubertal Development is Associated with Internalizing Symptoms through Family Conflict Across Sexes

As hypothesized in our theoretical model, physical pubertal development was positively associated with internalizing symptoms both directly and indirectly—through family conflict—across sexes, implying that family conflict may amplify pubertal effects on internalizing symptoms. These findings align with previous research suggesting that harsh parenting exacerbates pubertal timing effects on internalizing symptoms in females [37]. Our results thus extend previous findings by establishing indirect effects in both sexes, as early as ages 9-10, generalizing effects to pubertal development rather than pubertal timing, and more generally regarding family conflict rather than harsh parenting specifically.

Contrary to our theoretical model, associations between pubertal development, family conflict, and internalizing symptoms were not moderated by sex. At ages 9-10 in the ABCD sample, family conflict accounted for internalizing symptoms similarly in males and females, consistent with evidence suggesting that sex differences in family effects on internalizing symptoms do not emerge until mid-adolescence [63]. Furthermore, FES scores in our sample were lower in females, suggesting less family conflict. This aligns with females typically reporting greater family cohesion than males [64], which may heighten their long-term sensitivity to conflict and trigger internalizing symptoms at later pubertal stages [65], as family conflict increases throughout adolescence [66]. Therefore, the strength of the association between pubertal development and internalizing symptoms through family conflict is expected to increase over time, especially in females, and may contribute to the emergence of sex differences in these effects.

### 4.3. Endocrine and Neurocognitive Effects on Internalizing Symptoms are not yet Observable at Ages 9-10

Further contrary to our hypothesized theoretical model, our findings do not indicate effects of sex hormone levels and family conflict on internalizing symptoms through amygdala response to fearful faces. The ABCD sample at ages 9-10 may thus be too young to show the mechanisms proposed in our theoretical model, including associated sex differences, as the trend of greater internalizing symptoms in females is not yet observable in this sample. In fact, no sex differences in amygdala response to fearful faces were detected either, an effect consistently reported in adults [67]. Furthermore, no associations were found between amygdala response to fearful faces and internalizing symptoms, common to all unsupported hypotheses in our study attempting to establish brain function as the pathway from risk factors to internalizing symptoms [68]. Substantial research suggests that heightened amygdala response to fearful faces in adult females, but not males, may contribute to females’ greater vulnerability to internalizing disorders [69]. This association has however not yet been established in youth, consistent with our null findings. Previous research on adolescent depression also found no sex differences in amygdala response to negative stimuli at ages 13-17 [70]. Therefore, sex differences in amygdala reactivity to negative stimuli may only emerge in late-adolescence/early-adulthood as a potential risk factor for depression in females, and our sample may be too young to capture this effect.

It may also be too early, at ages 9-10, to observe direct effects of hormone levels on internalizing symptoms. Our findings did indicate higher levels of DHEA and testosterone in females relative to males, consistent with previous research suggesting that females begin puberty before males [71]. However, sex differences in hormone levels steadily increase throughout adolescence and endocrine effects on internalizing symptoms may thus take some time to emerge. Menarche has also been associated with the development of internalizing symptoms [72], particularly in females with heightened neurological vulnerability to fluctuating ovarian hormone levels [73]. However, these effects could not be reliably assessed in our sample given the small proportion of females reporting menarche. Nevertheless, as this was the first study exploring endocrine effects at such a young age, our null findings are still valuable in suggesting that it may be too early, at ages 9-10, to observe endocrine effects on internalizing symptoms.

### 4.3. Methodological Considerations

#### 4.3.1. Small Effect Sizes

Although we found some statistically significant results, effect sizes were small by traditional standards, e.g., *R*^2^ = .03% for the effect of family conflict on physical pubertal development. Such small effects may have multiple explanations. Firstly, research conducted on large samples [74], including the ABCD sample [75], has recently shown similarly small effect sizes, for example the statistically significant correlations between parental acceptance and total psychiatric problems (*r* = -.09) or between caffeine consumption and sleep problems (*r* = .04) [76]. Established benchmarks for interpreting effect sizes may thus be historically based on underpowered or biased studies overestimating effect sizes, and should therefore not be used to assess the meaningfulness of our effects [77]. Secondly, large-scale research initiatives are powered to detect small effects, which are suggested to more closely approximate true population values [78]. Small effect sizes in large, multisite studies of heterogeneous samples may also be due to increased statistical noise diluting effect sizes [79]. Nevertheless, large and diverse samples, such as the ABCD cohort, directly address the issue of low power and bias in research [80], and are thus important to overcome the typically low reproducibility of findings in psychology and neuroscience [81]. Thirdly, given that our sample is not clinical, we are likely looking at subtle variation in internalizing pathology within the normal range and would not expect to observe large effect sizes in this context and at this age. However, in developmental psychopathology, effects are expected to accumulate and grow over time [82]. Small effects like ours may thus additively explain a substantial proportion of variance in neurodevelopmental trajectories [83].

#### 4.3.2. Methodological Limitations

Although this study has valuable strengths, some methodological limitations should be acknowledged. Firstly, the fMRI emotional face n-back task may not have adequately captured emotional reactivity due to potential interference from the tasks’ working memory component. Motivated to perform well and focusing on evaluating whether stimuli matched those presented two trials back, subjects may have not paid explicit attention to the emotional valence of the presented stimuli. In fact, the construction of conscious emotional experience is argued to require the integration of both implicit (e.g., visceral) and explicit (e.g., attentional) emotional processes [84]. The emotional face n-back task may thus not be the most suitable measure of emotional reactivity. Nevertheless, our data shows evidence that the task did yield differential amygdala activations in the contrast between fearful and neutral faces, supporting its intended use to establish the neurocognitive effects depicted by our theoretical model.

Secondly, as our analyses were conducted on data collected at a single timepoint, no inferences on the directionality of direct and indirect effects could be made. Mediation analysis indeed requires longitudinal data to establish directionality [85]. However, mediation models tested with single timepoint data have shown acceptable performance in a recent study using simulations to compare different mediation models including an equal amount of data but varying numbers of timepoints [86]. Therefore, mediation analysis still served our aims of gaining a preliminary mechanistic understanding of how multilevel data may interact to yield internalizing symptoms through direct and indirect effects.

### 4.4. Implications and Future Directions

This is the first study taking a developmental perspective to concurrently investigate endocrine, neurocognitive, and psychosocial factors associated with emerging sex differences in internalizing symptoms in a 9-to 10-year-old sample. By taking an integrative approach, we considered the additive contributions of heterogenous risk factors on internalizing psychopathology and how they may interact with one another.

Our findings have theoretical implications in suggesting that sex differences in internalizing symptoms are not yet present among 9-to 10-year-olds. Considering the ABCD sample’s size and diversity, these findings valuably contribute to existing mixed research on the age at which sex differences emerge. Furthermore, null findings for direct and indirect effects of sex hormones and family conflict on internalizing symptoms through brain function suggest that it may also be too early, at ages 9-10, to observe interactive effects of endocrine, neurocognitive, and psychosocial risk factors.

In the field of developmental psychopathology, null findings serve as an important baseline for future longitudinal research [87]. Given that adolescence is a transitional developmental period, the presently hypothesized effects and associated sex differences are expected to emerge and grow over time [88]. As the ABCD study aims to prospectively follow its baseline cohort for ten years, it will provide an unprecedented opportunity to build on the present findings by testing our proposed theoretical model at different ages. Longitudinal investigation of the developmental trajectories of sex differences in internalizing symptoms will further allow us to assess the directionality of effects.

Future research on emerging sex differences in internalizing symptoms should also investigate fronto-limbic network connectivity rather than amygdala reactivity alone, given recent findings identifying sex differences in associations between amygdala functional connectivity and internalizing symptoms in neurotypical youth [89]. Furthermore, psychosocial effects of family and peer relationships need to be considered together, as youth dynamically shift from one to the other over the course of adolescence [90]. The longitudinal investigation of our theoretical model with the proposed additional considerations may have major clinical implications for the early diagnosis and detection of youth at risk by informing more targeted interventions for prevention and treatment.

## 5. Conclusion

Taking a developmental and multidimensional approach is necessary to understand the greater prevalence of internalizing symptoms in females. We concurrently assessed contributions from endocrine, neurocognitive, and psychosocial perspectives to study interacting mechanisms underlying emerging sex differences in a large and diverse sample of 9- to 10-year-olds. Consistent with previous research, effects of pubertal development on internalizing symptoms through family conflict were observed across sexes. However, it seems too early at this age to observe the surge of internalizing symptoms in females, as well as interactive effects of endocrine, neurocognitive, and psychosocial factors. By examining mechanisms of risk prior to the peak period for the onset of psychopathology, our findings provide an important baseline for future longitudinal research on the emergence of sex differences in internalizing symptoms.

## Acknowledgements & Funding

Data used in the preparation of this article were obtained from the Adolescent Brain Cognitive Development (ABCD) Study (https://abcdstudy.org), held in the NIMH Data Archive (NDA). This is a multisite, longitudinal study designed to recruit more than 10,000 children age 9-10 and follow them over 10 years into early adulthood. The ABCD Study is supported by the National Institutes of Health and additional federal partners under award numbers U01DA041048, U01DA050989, U01DA051016, U01DA041022, U01DA051018, U01DA051037, U01DA050987, U01DA041174, U01DA041106, U01DA041117, U01DA041028, U01DA041134, U01DA050988, U01DA051039, U01DA041156, U01DA041025, U01DA041120, U01DA051038, U01DA041148, U01DA041093, U01DA041089, U24DA041123, U24DA041147. A full list of supporters is available at https://abcdstudy.org/federal-partners.html. A listing of participating sites and a complete listing of the study investigators can be found at https://abcdstudy.org/consortium_members/. ABCD consortium investigators designed and implemented the study and/or provided data but did not necessarily participate in the analysis or writing of this report. This manuscript reflects the views of the authors and may not reflect the opinions or views of the NIH or ABCD consortium investigators. The ABCD data repository grows and changes over time. The ABCD data used in this report came from NDA Release 3.0 (DOI: 10.15154/1519007). DOIs can be found at https://dx.doi.org/10.15154/1519007.

The involvement of authors Robert Kohler and Sarah D Lichenstein was supported by National Institute on Drug Abuse grants T32DA007238 and K08DA051667 respectively.

## CRediT authorship contribution statement

**Bianca Serio:** Conceptualization; Data curation; Formal analysis; Investigation; Methodology; Software; Validation; Visualization; Roles/Writing - original draft; Writing - review & editing. **Robert Kohler:** Conceptualization; Data curation; Formal analysis; Investigation; Methodology; Software; Validation; Writing - review & editing. **Fengdan Ye**: Data Curation; Writing – Review & Editing. **Sarah D Lichenstein:** Conceptualization; Data Curation; Validation; Writing – Review & Editing. **Sarah W Yip**: Conceptualization; Data curation; Investigation; Methodology; Project administration; Resources; Supervision; Validation; Writing - review & editing.

## Data Availability

Data for this study is available through the ABCD NDA (https://nda.nih.gov/abcd).

## Appendix. Supplementary Material

Supplementary results associated with this article are appended.

## References

1. Achenbach, T.M., The classification of children’s psychiatric symptoms: A factor-analytic study. Psychological Monographs: General and applied, 1966. 80(7): p. 1.

2. Quay, H.C., Classification, in Psychopathological Disorders of Childhood, H.C. Quay and J.S. Werry, Editors. 1986, Wiley: New York. p. 1–34.

3. Eaton, N.R., et al., An invariant dimensional liability model of gender differences in mental disorder prevalence: Evidence from a national sample. Journal of abnormal psychology, 2012. 121(1): p. 282.

4. Mendle, J., Why puberty matters for psychopathology. Child Development Perspectives, 2014. 8(4): p. 218–222.

5. Kessler, R.C., et al., Sex and depression in the National Comorbidity Survey. II: Cohort effects. Journal of affective disorders, 1994. 30(1): p. 15–26.

6. Breslau, J., et al., Sex differences in recent first-onset depression in an epidemiological sample of adolescents. Translational psychiatry, 2017. 7(5): p. e1139–e1139.

7. Letcher, P., et al., Precursors and correlates of anxiety trajectories from late childhood to late adolescence. Journal of Clinical Child & Adolescent Psychology, 2012. 41(4): p. 417–432.

8. Sweeting, H. and P. West, Sex differences in health at ages 11, 13 and 15. Social science & medicine, 2003. 56(1): p. 31–39.

9. Wade, T.J., J. Cairney, and D.J. Pevalin, Emergence of gender differences in depression during adolescence: National panel results from three countries. Journal of the American Academy of Child & Adolescent Psychiatry, 2002. 41(2): p. 190–198.

10. Soares, C.N. and B. Zitek, Reproductive hormone sensitivity and risk for depression across the female life cycle: A continuum of vulnerability? Journal of psychiatry & neuroscience, 2008. 33(4): p. 331–343.

11. Lewis, G., et al., The association between pubertal status and depressive symptoms and diagnoses in adolescent females: A population-based cohort study. PloS one, 2018. 13(6): p. e0198804.

12. Reardon, L.E., E.W. Leen-Feldner, and C. Hayward, A critical review of the empirical literature on the relation between anxiety and puberty. Clinical psychology review, 2009. 29(1): p. 1–23.

13. Mendle, J., et al., Understanding puberty and its measurement: ideas for research in a new generation. Journal of Research on Adolescence, 2019. 29(1): p. 82–95.

14. Han, G., et al., Adolescents’ internalizing and externalizing problems predict their affect-specific HPA and HPG axes reactivity. Developmental Psychobiology, 2015. 57(6): p. 769–785.

15. Mulligan, E.M., et al., Increased dehydroepiandrosterone (DHEA) is associated with anxiety in adolescent girls. Psychoneuroendocrinology, 2020. 119: p. 104751.

16. Chronister, B.N., et al., The Effects of Testosterone, Estradiol, Dehydroepiandrosterone, and Cortisol on Anxiety and Depression Symptoms in Ecuadorian Adolescents. American Academy of Pediatrics, 2021. 147(3).

17. Copeland, W.E., et al., Early pubertal timing and testosterone associated with higher levels of adolescent depression in girls. Journal of the American Academy of Child & Adolescent Psychiatry, 2019. 58(12): p. 1197–1206.

18. Angold, A., et al., Pubertal changes in hormone levels and depression in girls. Psychological medicine, 1999. 29(5): p. 1043–1053.

19. Rubinow, D.R. and P.J. Schmidt, Sex differences and the neurobiology of affective disorders. Neuropsychopharmacology, 2019. 44(1): p. 111–128.

20. Borrow, A. and R.J. Handa, Estrogen receptors modulation of anxiety-like behavior. Vitamins and hormones, 2017. 103: p. 27–52.

21. Sripada, R.K., et al., DHEA enhances emotion regulation neurocircuits and modulates memory for emotional stimuli. Neuropsychopharmacology, 2013. 38(9): p. 1798–1807.

22. Klein, S., et al., Increased neural reactivity to emotional pictures in men with high hair testosterone concentrations. Social cognitive and affective neuroscience, 2019. 14(9): p. 1009–1016.

23. Henningsson, S., et al., Role of emotional processing in depressive responses to sex-hormone manipulation: A pharmacological fMRI study. Translational psychiatry, 2015. 5(12): p. e688–e688.

24. Guyer, A.E., et al., A developmental examination of amygdala response to facial expressions. Journal of Cognitive neuroscience, 2008. 20(9): p. 1565–1582.

25. Guyer, A.E., J.S. Silk, and E.E. Nelson, The neurobiology of the emotional adolescent: From the inside out. Neuroscience & Biobehavioral Reviews, 2016. 70: p. 74–85.

26. Jiang, N., et al., Negative parenting affects adolescent internalizing symptoms through alterations in amygdala-prefrontal circuitry: A longitudinal twin study. Biological Psychiatry, 2021. 89(6): p. 560–569.

27. Callaghan, B.L., et al., Amygdala resting connectivity mediates association between maternal aggression and adolescent major depression: a 7-year longitudinal study. Journal of the American Academy of Child & Adolescent Psychiatry, 2017. 56(11): p. 983–991.

28. Hamlat, E.J., et al., Early pubertal timing predicts onset and recurrence of depressive episodes in boys and girls. Journal of Child Psychology and Psychiatry, 2020. 61(11): p. 1266–1274.

29. Graber, J.A., et al., Is pubertal timing associated with psychopathology in young adulthood? Journal of the American Academy of Child & Adolescent Psychiatry, 2004. 43(6): p. 718–726.

30. Negriff, S. and E.J. Susman, Pubertal timing, depression, and externalizing problems: A framework, review, and examination of gender differences. Journal of Research on Adolescence, 2011. 21(3): p. 717–746.

31. Barendse, M., et al., Multi-method assessment of pubertal timing and associations with internalizing psychopathology in early adolescent girls. PsyArXiv Preprints, 2020.

32. Mouritsen, A., et al., The pubertal transition in 179 healthy Danish children: Associations between pubarche, adrenarche, gonadarche, and body composition. European journal of endocrinology, 2013. 168(2): p. 129–136.

33. Krauss, S., U. Orth, and R.W. Robins, Family environment and self-esteem development: A longitudinal study from age 10 to 16. Journal of Personality and Social Psychology, 2020. 119(2): p. 457.

34. Jiang, Y., et al., Buffering the effects of peer victimization on adolescent non-suicidal self-injury: The role of self-compassion and family cohesion. Journal of Adolescence, 2016. 53: p. 107–115.

35. Compian, L.J., L.K. Gowen, and C. Hayward, The interactive effects of puberty and peer victimization on weight concerns and depression symptoms among early adolescent girls. The Journal of Early Adolescence, 2009. 29(3): p. 357–375.

36. Stadler, C., et al., Peer-victimization and mental health problems in adolescents: Are parental and school support protective? Child Psychiatry & Human Development, 2010. 41(4): p. 371–386.

37. Deardorff, J., et al., Pubertal timing and Mexican-origin girls’ internalizing and externalizing symptoms: The influence of harsh parenting. Developmental psychology, 2013. 49(9): p. 1790.

38. Pfeifer, J.H. and N.B. Allen, Puberty initiates cascading relationships between neurodevelopmental, social, and internalizing processes across adolescence. Biological Psychiatry, 2020. 89(2): p. 99–108.

39. Jernigan, T.L., S.A. Brown, and G.J. Dowling, The adolescent brain cognitive development study. Journal of research on adolescence: the official journal of the Society for Research on Adolescence, 2018. 28(1): p. 154.

40. Barch, D.M., et al., Demographic, physical and mental health assessments in the adolescent brain and cognitive development study: Rationale and description. Developmental Cognitive Neuroscience, 2018. 32: p. 55–66.

41. Volkow, N.D., et al., The conception of the ABCD study: From substance use to a broad NIH collaboration. Developmental cognitive neuroscience, 2018. 32: p. 4–7.

42. Garavan, H., et al., Recruiting the ABCD sample: Design considerations and procedures. Developmental cognitive neuroscience, 2018. 32: p. 16–22.

43. Heeringa, S.G. and P.A. Berglund, A guide for population-based analysis of the Adolescent Brain Cognitive Development (ABCD) Study baseline data. BioRxiv, 2020.

44. Paul, S.E., et al., Associations between prenatal cannabis exposure and childhood outcomes: Results from the ABCD study. JAMA Psychiatry, 2021. 78(1): p. 64–76.

45. Achenbach, T.M. and C. Edelbrock, Child behavior checklist. Burlington (Vt), 1991. 7: p. 371–392.

46. Uban, K.A., et al., Biospecimens and the ABCD study: Rationale, methods of collection, measurement and early data. Developmental cognitive neuroscience, 2018. 32: p. 97–106.

47. Herting, M.M., et al., Correspondence between perceived pubertal development and hormone levels in 9-10 year-olds from the Adolescent Brain Cognitive Development study. Frontiers in Endocrinology, 2020. 11.

48. Petersen, A.C., et al., A self-report measure of pubertal status: Reliability, validity, and initial norms. Journal of youth and adolescence, 1988. 17(2): p. 117–133.

49. Carskadon, M.A. and C. Acebo, A self-administered rating scale for pubertal development. Journal of Adolescent Health, 1993. 14(3): p. 190–195.

50. Moos, R.H. and B. Humphrey, Preliminary manual for family environment scale, work environment scale, group environment scale. 1974, Palo Alto, CA:Consulting Psychologists Press.

51. Casey, B., et al., The adolescent brain cognitive development (ABCD) study: Imaging acquisition across 21 sites. Developmental cognitive neuroscience, 2018. 32: p. 43–54.

52. Cohen, A., et al., The impact of emotional cues on short-term and long-term memory during adolescence. Program No. 90.25 Neuroscience Meeting Planner. 2016, San Diego, CA: Society for Neuroscience.

53. Kerestes, R., et al., Abnormal prefrontal activity subserving attentional control of emotion in remitted depressed patients during a working memory task with emotional distracters. Psychological medicine, 2012. 42(1): p. 29.

54. Conley, M.I., et al., The racially diverse affective expression (RADIATE) face stimulus set. Psychiatry research, 2018. 270: p. 1059–1067.

55. Tottenham, N., et al., The NimStim set of facial expressions: Judgments from untrained research participants. Psychiatry research, 2009. 168(3): p. 242–249.

56. Cureton, E.E., Rank-biserial correlation. Psychometrika, 1956. 21(3): p. 287–290.

57. Tingley, D., et al., Mediation: R package for causal mediation analysis. Journal of Statistical Software, 2014. 59(5).

58. MacKinnon, D.P., A.J. Fairchild, and M.S. Fritz, Mediation analysis. Annual Review of Psychology, 2007. 58: p. 593–614.

59. Dick, A.S., et al., Meaningful associations in the adolescent brain cognitive development study. NeuroImage, 2021: p. 118262.

60. Efron, B., Better bootstrap confidence intervals. Journal of the American statistical Association, 1987. 82(397): p. 171–185.

61. Preacher, K.J., D.D. Rucker, and A.F. Hayes, Addressing moderated mediation hypotheses: Theory, methods, and prescriptions. Multivariate behavioral research, 2007. 42(1): p. 185–227.

62. Ruchkin, V. and M. Schwab-Stone, A longitudinal study of somatic complaints in urban adolescents: The role of internalizing psychopathology and somatic anxiety. Journal of youth and adolescence, 2014. 43(5): p. 834–845.

63. Crawford, T.N., et al., Internalizing symptoms in adolescents: Gender differences in vulnerability to parental distress and discord. Journal of research on adolescence, 2001. 11(1): p. 95–118.

64. Sze, T.-M., et al., Sex differences in the development of perceived family cohesion and depressive symptoms in Taiwanese adolescents. Psychological reports, 2013. 113(1): p. 54–72.

65. Lewis, A., et al., Gender differences in adolescent depression: Differential female susceptibility to stressors affecting family functioning. Australian Journal of Psychology, 2015. 67(3): p. 131–139.

66. Steinberg, L., Reciprocal relation between parent-child distance and pubertal maturation. Developmental psychology, 1988. 24(1): p. 122.

67. Andreano, J.M., B.C. Dickerson, and L.F. Barrett, Sex differences in the persistence of the amygdala response to negative material. Social Cognitive and Affective Neuroscience, 2014. 9(9): p. 1388–1394.

68. Drevets, W.C., J.L. Price, and M.L. Furey, Brain structural and functional abnormalities in mood disorders: Implications for neurocircuitry models of depression. Brain structure and function, 2008. 213(1-2): p. 93–118.

69. Dickie, E.W. and J.L. Armony, Amygdala responses to unattended fearful faces: Interaction between sex and trait anxiety. Psychiatry Research: Neuroimaging, 2008. 162(1): p. 51–57.

70. Yang, T.T., et al., Adolescents with major depression demonstrate increased amygdala activation. Journal of the American Academy of Child & Adolescent Psychiatry, 2010. 49(1): p. 42–51.

71. Marshall, W.A., Puberty, in Human growth, F. Falkner and J.M. Tanner, Editors. 1978, Springer: New York. p. 141–181.

72. Mendle, J., R.M. Ryan, and K.M. McKone, Age at menarche, depression, and antisocial behavior in adulthood. Pediatrics, 2018. 141(1).

73. Barth, C., A. Villringer, and J. Sacher, Sex hormones affect neurotransmitters and shape the adult female brain during hormonal transition periods. Frontiers in neuroscience, 2015. 9: p. 37.

74. Miller, K.L., et al., Multimodal population brain imaging in the UK Biobank prospective epidemiological study. Nature neuroscience, 2016. 19(11): p. 1523–1536.

75. Marek, S., et al., Towards reproducible brain-wide association studies. BioRxiv, 2020.

76. Owens, M.M., et al., Recalibrating expectations about effect size: A multi-method survey of effect sizes in the ABCD study. PLoS ONE, 2021. 16(9): p. e0257535.

77. Schäfer, T. and M.A. Schwarz, The meaningfulness of effect sizes in psychological research: Differences between sub-disciplines and the impact of potential biases. Frontiers in Psychology, 2019. 10: p. 813.

78. Dick, A.S., et al., Meaningful associations in the adolescent brain cognitive development study. NeuroImage, 2021. 239: p. 118262.

79. Feaster, D.J., S. Mikulich-Gilbertson, and A.M. Brincks, Modeling site effects in the design and analysis of multi-site trials. The American journal of drug and alcohol abuse, 2011. 37(5): p. 383–391.

80. Ioannidis, J.P., Why most published research findings are false. PLoS medicine, 2005. 2(8): p. e124.

81. Button, K.S., et al., Power failure: why small sample size undermines the reliability of neuroscience. Nature reviews neuroscience, 2013. 14(5): p. 365–376.

82. Wenar, C. and P. Kerig, Developmental psychopathology: From infancy through adolescence. 4th ed. 2000, McGraw-Hill.

83. Boyle, E.A., Y.I. Li, and J.K. Pritchard, An expanded view of complex traits: from polygenic to omnigenic. Cell, 2017. 169(7): p. 1177–1186.

84. Quirin, M. and R.D. Lane, The construction of emotional experience requires the integration of implicit and explicit emotional processes. Behavioral and Brain Sciences, 2012. 35(3): p. 159.

85. Maxwell, S.E. and D.A. Cole, Bias in cross-sectional analyses of longitudinal mediation. Psychological methods, 2007. 12(1): p. 23.

86. Cain, M.K., Z. Zhang, and C. Bergeman, Time and other considerations in mediation design. Educational and psychological measurement, 2018. 78(6): p. 952–972.

87. Luna, B., B. Tervo-Clemmens, and F.J. Calabro, Considerations when characterizing adolescent neurocognitive development. Biological psychiatry, 2021. 89(2): p. 96–98.

88. Casey, B., S. Getz, and A. Galvan, The adolescent brain. Developmental review, 2008. 28(1): p. 62–77.

89. Padgaonkar, N., et al., Sex differences in internalizing symptoms and amygdala functional connectivity in neurotypical youth. Developmental cognitive neuroscience, 2020. 44: p. 100797.

90. Larson, R.W., et al., Changes in adolescents’ daily interactions with their families from ages 10 to 18: Disengagement and transformation. Developmental psychology, 1996. 32(4): p. 744.

