## Supplementary Material for "A Multidimensional Approach to Understanding the Emergence of Sex Differences in Internalizing Symptoms in Adolescence"

***Supplementary Methods***

Recommended quality control inclusion criteria provided in ABCD Release 3.0 for the imaging data

nBack tfMRI series passed the raw quality control; T1 series passed the raw quality control; no behavioral performance flags were raised (i.e., < 60% 0-back or 2-back accuracy); degrees of freedom > 200; total number of trials = 100; E-prime timing was matched or E-prime mismatch was recommended to be ignored; fMRI B0 unwarp was available; FreeSurfer quality control was passed; fMRI manual post-processing quality control was passed; fMRI registration to T1w < 19; fMRI maximum dorsal cutoff score < 65; and fMRI maximum ventral cutoff score < 60.

### Imaging Procedure

A neuroimaging protocol was followed by all ABCD sites and harmonized for three 3T scanner types (Philips, Siemens Prisma, and General Electric 750) [1]. Multi-channel coils were used for multiband echo planar imaging acquisitions. Stimulus presentation for the fMRI emotional face n-back task was programmed in E-Prime Professional 2.0. Imaging parameters are reported by Casey et al. [1].

The ABCD study’s Data Analysis and Informatics Center was responsible for preprocessing all imaging data with the Multi-Modal Processing Stream (MMPS), a software package developed for the ABCD study specifically. The MMPS includes primarily MATLAB functions and relies on publicly available neuroimaging software packages such as Analysis of Functional NeuroImages (AFNI), FreeSurfer, and FMRIB Software Library (FSL). AFNI’s 3dvolreg was used to register each frame to the first in order to correct for head motion [2]. The ABCD study’s preprocessing pipeline [3] is publicly available as an independently executable and self-contained platform at <https://www.nitrc.org/projects/abcd_study>. A general linear model was implemented in AFNI 3dDeconvolve to compute changes in the blood-oxygen-level-dependent (BOLD) signal related to the fMRI emotional face n-back task. FreeSurfer’s subcortical segmentation [4] was used to for the parcellation of the amygdala as our region of interest (ROI).

***Supplementary Results***

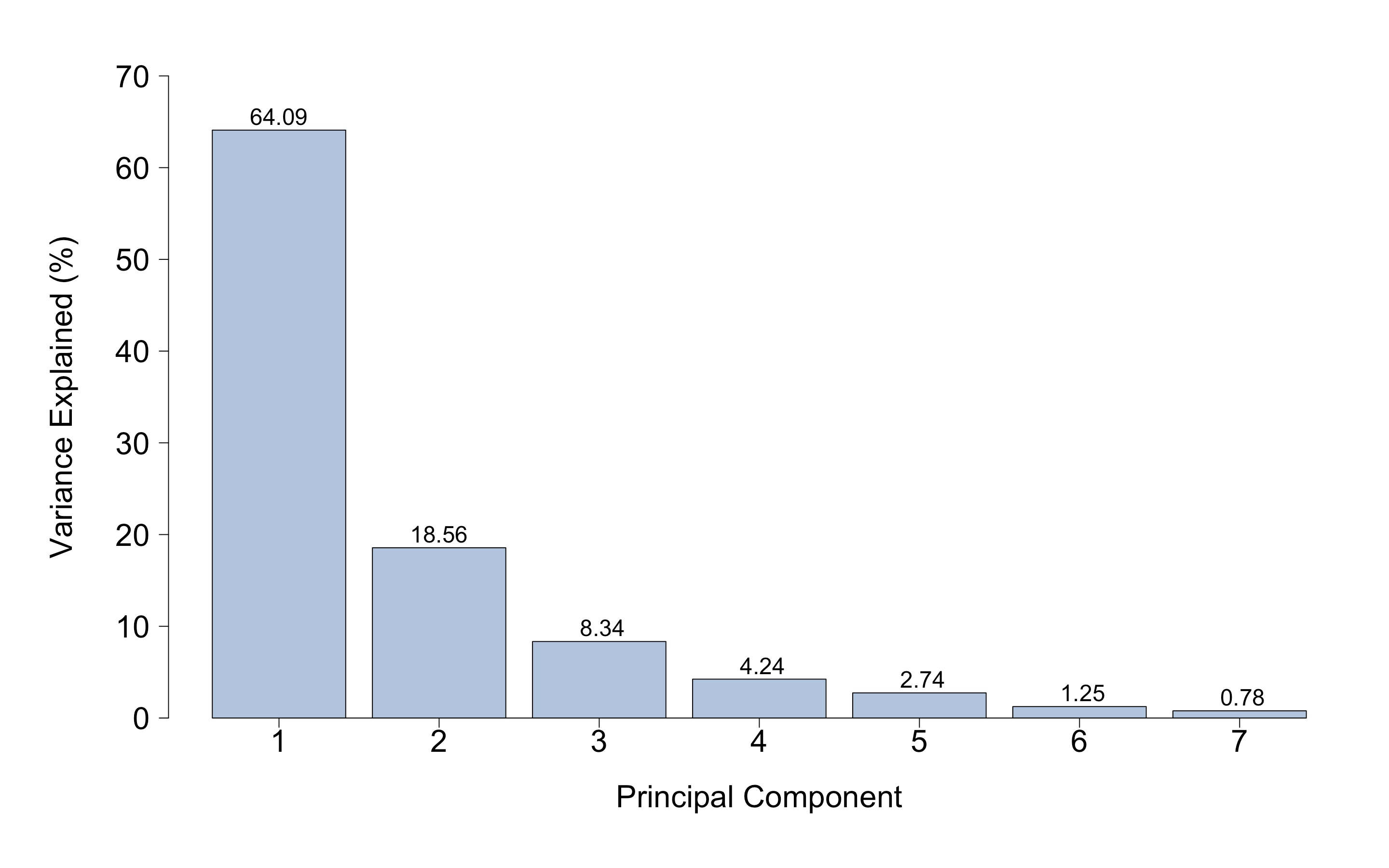

*Figure S1.* Variance explained by each principal component obtained with Principal Component Analysis

| **Factors included in PCA (*n* = 11,870)** |  | **PC1** | **PC2** | **PC3** | **PC4** | **PC5** | **PC6** | **PC7** |
| --- | --- | --- | --- | --- | --- | --- | --- | --- |
| Anxious/Depressed |  | 0.415 | 0.224 | -0.392 | 0.012 | -0.459 | 0.515 | 0.385 |
| Withdrawn/Depressed |  | 0.353 | 0.206 | 0.704 | -0.554 | -0.172 | -0.002 | 0.031 |
| Somatic |  | 0.338 | -0.591 | -0.007 | 0.004 | 0.060 | -0.416 | 0.601 |
| DSM Depression |  | 0.398 | 0.146 | 0.346 | 0.788 | -0.162 | -0.155 | -0.173 |
| DSM Anxiety |  | 0.411 | 0.196 | -0.477 | -0.258 | -0.109 | -0.559 | -0.417 |
| DSM Somatic |  | 0.291 | -0.673 | 0.006 | -0.072 | -0.054 | 0.414 | -0.533 |
| Stress |  | 0.421 | 0.216 | -0.064 | 0.007 | 0.846 | 0.233 | 0.052 |

*Table S2.* Principal Component Analysis factor loadings. PCA, principal component analysis; PC, principal component; n, number of subjects; DSM, Diagnostic and Statistical Manual of Mental Disorders. Positive and negative loadings respectively indicate positive and negative correlations between PCs and factors.

| **Hypothesis** | **Model** | **Predictor** | **β** |  | ***df*** |  | ***t*** |  | ***p*** |
| --- | --- | --- | --- | --- | --- | --- | --- | --- | --- |
| Direct and indirect effects of DHEA levels on internalizing symptoms through amygdala response to fearful faces | 1 | DHEA | 0.003 |  | 6,954 |  | 0.19 |  | .851 |
|  |  | Sex (M) | -0.029 |  | 6,720 |  | -1.23 |  | .220 |
|  |  | Collection duration | 0.008 |  | 7,092 |  | 0.65 |  | .516 |
|  |  | DHEA*Sex (M) | -0.011 |  | 7,028 |  | -0.43 |  | .668 |
|  |  | DHEA*Collection duration | -0.027 |  | 7,080 |  | -1.38 |  | .168 |
|  | 2 | DHEA | 0.021 |  | 6,934 |  | 1.41 |  | .158 |
|  |  | Sex (M) | 0.127 |  | 7,072 |  | 5.33 |  | **.000*** |
|  |  | AMY | -0.014 |  | 6,538 |  | -0.84 |  | .404 |
|  |  | Collection duration | 0.004 |  | 6,883 |  | 0.38 |  | .708 |
|  |  | DHEA*Sex (M) | -0.002 |  | 6,893 |  | 0.09 |  | .927 |
|  |  | AMY*Sex (M) | 0.017 |  | 6,656 |  | 0.73 |  | .468 |
|  |  | DHEA*Collection duration | -0.013 |  | 7,089 |  | -0.69 |  | .491 |
| Direct and indirect effects of testosterone levels on internalizing symptoms through amygdala response to fearful faces | 1 | TEST | 0.006 |  | 7,046 |  | 0.32 |  | .746 |
|  |  | Sex (M) | -0.031 |  | 6,746 |  | -1.29 |  | .197 |
|  |  | Collection duration | 0.006 |  | 7,114 |  | 0.30 |  | .620 |
|  |  | Time of the day | 0.016 |  | 6,555 |  | 1.30 |  | .194 |
|  |  | TEST*Sex (M) | -0.021 |  | 6,889 |  | -0.89 |  | .376 |
|  |  | TEST*Collection duration | -0.018 |  | 7,137 |  | -0.94 |  | .350 |
|  |  | TEST*Time of the day | 0.031 |  | 7,121 |  | 2.53 |  | .011 |
|  | 2 | TEST | 0.011 |  | 6,951 |  | 0.63 |  | .526 |
|  |  | Sex (M) | 0.131 |  | 7,096 |  | 5.59 |  | **.000*** |
|  |  | AMY | -0.014 |  | 6,682 |  | -0.79 |  | .428 |
|  |  | Collection duration | 0.004 |  | 6,995 |  | 0.30 |  | .766 |
|  |  | Time of the day | 0.024 |  | 4,751 |  | 1.81 |  | .070 |
|  |  | TEST*Sex (M) | 0.027 |  | 6,911 |  | 1.15 |  | .250 |
|  |  | AMY*Sex (M) | 0.015 |  | 6,754 |  | 0.66 |  | .510 |
|  |  | TEST*Collection duration | -0.010 |  | 7,105 |  | -0.56 |  | .574 |
|  |  | TEST*Time of the day | 0.001 |  | 6,332 |  | 0.06 |  | .953 |
| Direct and indirect effects of estradiol levels on internalizing symptoms through amygdala response to fearful faces | 1 | ESTR | 0.024 |  | 3,009 |  | 1.36 |  | .174 |
|  |  | Collection duration | 0.006 |  | 3,396 |  | 0.33 |  | .743 |
|  |  | Time of the day | 0.016 |  | 5,744 |  | 0.86 |  | .390 |
|  |  | Caffeine | 0.024 |  | 3,298 |  | 1.35 |  | .177 |
|  |  | ESTR*Collection duration | -0.004 |  | 3,391 |  | -0.16 |  | .877 |
|  |  | ESTR*Time of the day | 0.006 |  | 3,367 |  | 0.37 |  | .713 |
|  |  | ESTR*Caffeine | 0.001 |  | 3,391 |  | 0.07 |  | .947 |
|  | 2 | ESTR | 0.006 |  | 3,296 |  | 0.36 |  | .717 |
|  |  | AMY | -0.015 |  | 3,247 |  | -0.90 |  | .370 |
|  |  | Collection duration | 0.002 |  | 3,301 |  | 0.10 |  | .924 |
|  |  | Time of the day | 0.012 |  | 2,277 |  | 0.67 |  | .505 |
|  |  | Caffeine | 0.007 |  | 3,395 |  | 0.43 |  | .670 |
|  |  | ESTR*Collection duration | 0.001 |  | 3,317 |  | 0.02 |  | .984 |
|  |  | ESTR*Time of the day | 0.005 |  | 3,318 |  | 0.29 |  | .775 |
|  |  | ESTR*Caffeine | 0.019 |  | 3,383 |  | 1.37 |  | .172 |
| Direct and indirect effects of family conflict on internalizing symptoms through amygdala response to fearful faces | 1 | FES | 0.008 |  | 7,542 |  | 0.50 |  | .620 |
|  |  | Sex (M) | -0.024 |  | 7,014 |  | -1.04 |  | .301 |
|  |  | FES*Sex (M) | -0.008 |  | 7,690 |  | -0.34 |  | .732 |
|  | 2 | FES | 0.089 |  | 6,758 |  | 5.62 |  | **.000*** |
|  |  | Sex (M) | 0.107 |  | 7,663 |  | 4.75 |  | **.000*** |
|  |  | AMY | -0.024 |  | 6,833 |  | -1.44 |  | .151 |
|  |  | FES*Sex (M) | 0.002 |  | 6,742 |  | 0.07 |  | .943 |
|  |  | AMY*Sex (M) | 0.023 |  | 7,082 |  | 1.04 |  | .298 |

*Table S3.* Test statistics of the GLME models required to conduct mediation analyses for Hypotheses 1 and 2. GLME, generalized linear mixed-effect; β, Standardized beta coefficient; *df*, degrees of freedom; DHEA, dehydroepiandrosterone; TEST, testosterone; ESTR, estradiol; FES, Family Environment Scale (score); AMY, amygdala response to fearful faces; M, male. Bold and * denotes statistical significance at *p* < .001. Model 1 predicts amygdala response to fearful faces, and Model 2 predicts internalizing symptoms (PC1 scores).

**References**

1.	Casey, B., et al., *The adolescent brain cognitive development (ABCD) study: Imaging acquisition across 21 sites.* Developmental cognitive neuroscience, 2018. **32**: p. 43-54.

2.	Cox, R.W., *AFNI: Software for analysis and visualization of functional magnetic resonance neuroimages.* Computers and Biomedical research, 1996. **29**(3): p. 162-173.

3.	Hagler Jr, D.J., et al., *Image processing and analysis methods for the Adolescent Brain Cognitive Development Study.* Neuroimage, 2019. **202**: p. 116091.

4.	Fischl, B., et al., *Whole brain segmentation: Automated labeling of neuroanatomical structures in the human brain.* Neuron, 2002. **33**(3): p. 341-355.
